# OTX2 controls chromatin accessibility to direct somatic versus germline differentiation

**DOI:** 10.1101/2025.02.18.638827

**Authors:** Elisa Barbieri, Ian Chambers

## Abstract

The choice between somatic and germline fates is essential for species survival. This choice occurs in embryonic epiblast cells, as these cells are competent for both somatic and germline differentiation. The transcription factor OTX2 regulates the choice between somatic and germline fates, as *Otx2*-null epiblast-like cells (EpiLCs) form primordial germ cell-like cells (PGCLCs) with enhanced efficiency. However, the mechanisms by which OTX2 achieves this function are not fully characterized. Here we show that OTX2 controls chromatin accessibility at specific chromatin loci to modulate gene transcription and enable somatic differentiation. By performing CUT&RUN for OTX2 and ATAC-seq in wild-type and *Otx2*-null embryonic stem cells and EpiLCs, we identified regions where OTX2 binding opens chromatin, either alone or in cooperation with a ZIC family protein. Enforced OTX2 expression maintains accessibility at these regions and, in addition, induces opening of ∼4,000 somatic-associated regions in cells differentiating in the presence of PGC-inducing cytokines. Once cells have acquired germline identity, these additional somatic-associated regions can no longer respond to OTX2 and remain closed. Our results indicate that OTX2 works in cells with dual competence for both somatic and germline differentiation to increase accessibility of somatic regulatory regions and induce the somatic fate at the expense of the germline.

## Introduction

During embryonic development, cells are faced with choices that determine their fate on multiple occasions. A critical step is the choice between somatic and germline differentiation which occurs early in mammalian development shorty after implantation. In mouse embryos at day 6.5, cells in the embryo proper express OTX2, a transcription factor associated with epiblast identity both *in vivo* and *in vitro* and with a critical function in neural development [1–5]. Cells in the posterior proximal region of the epiblast respond to external signals, in particular bone morphogenic factor (BMP) 4, secreted from the adjacent extraembryonic tissue [6,7]. At this time, while the rest of the epiblast continues to express OTX2, this group of cells downregulate OTX2. Subsequently, these cells go on to express primordial germ cell (PGC)- associated transcription factors BLIMP1, PRDM14 and AP2γ [7–14]. This process can be recapitulated *in vitro*. Naïve embryonic stem cells (ESCs) can be differentiated into PGC-like cells (PGCLCs) via a transient population of formative pluripotent cells called epiblast-like cells (EpiLCs). EpiLCs are able to respond to BMP4 and are therefore considered competent for germline development [15,16]. These cells downregulate OTX2 and subsequently induce expression of PGC-associated TFs [8,15].

The differential expression of OTX2, which is high in the epiblast but becomes repressed as cells enter the germline, is suggestive of a role for OTX2 in the choice between somatic and germline differentiation. Indeed*, in vivo*, embryos lacking *Otx2* show an increase in PGC numbers at embryonic day 7.5 [8]. In addition, *Otx2*-null ESCs can generate PGCLCs from EpiLCs with higher efficiency than wild-type cells [8].

Cells in the posterior epiblast, as well as EpiLCs *in vitro*, possess a dual competence for both somatic and germline differentiation. The choice of which fate to follow is based on several factors, among which the reactivation of the naïve gene regulatory network plays a pivotal role. OTX2 has been associated with the ability to repress the naïve gene regulatory network. Indeed, OTX2 antagonises the activity of NANOG in ESCs [17] and OTX2 binding sequences have been identified in the regulatory regions of *Nanog*, *Oct4* and *Sox2* [18]. Deletion of OTX2 binding elements in the *Nanog* and *Oct4* regulatory regions leads to the increased expression of *Nanog* and *Oct4* in EpiLCs and an increased yield of PGCLCs [19].

OTX2 not only represses naïve pluripotency. OTX2 overexpression in ESCs induces the exit from naïve pluripotency and the expression of formative and primed pluripotency associated genes [2]. This suggests that OTX2 can actively instruct the cells towards a more differentiated cell state. Moreover, the increase in OTX2 expression during the transition of ESCs to EpiLCs leads to the redistribution of OCT4 on chromatin, contributing to the establishment of the formative and primed gene regulatory network [20,21]. Together these observations suggest that OTX2 has a more prominent role in determining the somatic fate of cells with competence for both germline and somatic differentiation.

In this work we show that OTX2 is able to open chromatin at specific somatic regulatory regions, for example at the *Fgf5* enhancers, priming the cells towards the somatic fate. OTX2 induces chromatin accessibility early during the EpiLC to PGCLC transition, when cells possess dual competence, instructing cells towards the somatic fate at the expense of the germline.

## Results

### OTX2 localization at chromatin

To investigate how OTX2 acts on chromatin to determine differentiation choice, the chromatin binding profile of OTX2 was analyzed in cells competent for germline differentiation. Wild-type ESCs were differentiated into EpiLCs for 44 hours and Cleavage Under Targets &Release Using Nuclease (CUT&RUN [22]) was performed in both ESCs and EpiLCs (Figure 1A). A total of 7,136 OTX2-bound regions were identified (Figures 1B). The majority of OTX2-bound regions (4,443) are EpiLC-specific but 1,429 regions are ESC-specific and 1,264 regions are bound by OTX2 in both ESCs and EpiLCs. Heatmaps shows the distribution of OTX2 binding to these 7,136 regions (Figure 1C). While ESC-specific and EpiLC-specific regions had higher OTX2 occupancy in the respective cell type (Figures 1C-D), common regions showed higher occupancy by OTX2 in EpiLCs (Figures 1C), in line with the higher OTX2 protein expression in EpiLCs [20]. This was borne out in the analysis of the average signal of OTX2 binding to these regions which is also slightly higher in EpiLCs (Figure 1D). Indeed, common regions showed the highest average signal of all sites in both ESCs and EpiLCs, suggesting that although already present in ESCs, OTX2 binds strongly or more frequently to these regions in EpiLCs compared to ESCs (Figures 1D). Examples of ESC-specific, common and EpiLC-specific OTX2 peaks are enhancers of *Tet2, Mycn* and *Fgf5*, respectively. *Tet2* intragenic enhancer shows strong binding of OTX2 in wild-type ESCs but not in EpiLCs. OTX2 binds *Mycn* enhancer in both cell types with higher intensity in EpiLCs compared to ESCs. In contrast, the *Fgf5* downstream enhancers E1, E2 and E3 are bound by OTX2 only in EpiLCs (Figure 1E).

**Figure 1.**
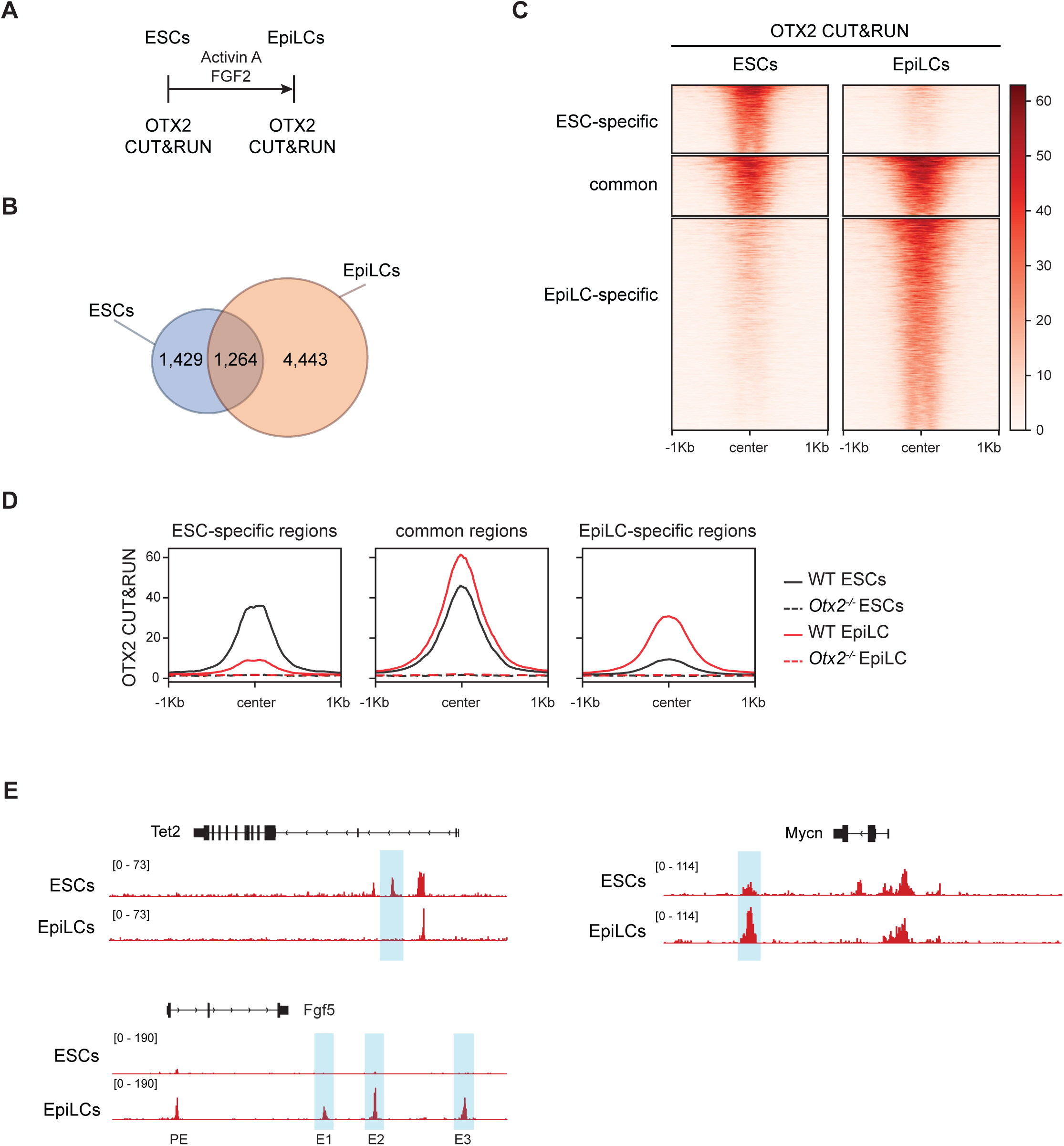
OTX2 chromatin binding during the ESC to EpiLC transition. A) Summary of OTX2 CUT&RUN samples analyzed in this study. B) Venn diagram of OTX2 bound regions in ESCs (blue) and EpiLCs (orange). C) Heatmap of OTX2 CUT&RUN signal, showing ESC-specific, EpiLC-specific and common regions. D) Read density profiles of OTX2 CUT&RUN in wild-type and *Otx2^-/-^*ESCs, wild-type and *Otx2^-/-^* EpiLCs at ESC-specific, common and EpiLC-specific OTX2-bound regions. E) OTX2 CUT&RUN tracks showing examples of ESC-specific (*Tet2*), common (*Mycn*) and EpiLC-specific (*Fgf5*) OTX2-bound regions. Nomenclature of *Fgf5* enhancers from [20].

OTX2-bound regions are located distal to the transcription start sites in both ESCs and EpiLCs (Figure S1A). Although the binding of OTX2 to promoters increases in common and EpiLC-specific regions compared to ESC-specific regions, the majority of OTX2 remains bound to distal regions (Figure S1A). This suggests that in all three cases OTX2 binds mostly to putative enhancers.

To investigate differences between the OTX2-bound sites, motif analysis on ESC-specific, common and EpiLC-specific OTX2 peaks was performed. As expected, the OTX2 motif was enriched in all subsets (Figure S1B). Together with OTX2-like motifs recognized by GSC and CRX, these were among the most enriched in all three subsets. Consistent with the known interaction of OCT4 with OTX2 [20], the OCT/SOX motif was also enriched in all three datasets (Figure S1B). Differences between the subsets emerge when analyzing other enriched motifs. ESC-specific and common regions are enriched for SOX and KLF family motifs and EpiLC-specific regions are enriched for ZIC family motifs (Figure S1B). Gene ontology analysis of the genes closest to OTX2-bound regions, and that are therefore likely to be targets of OTX2, reveals that both ESC-specific and EpiLC-specific regions associate with genes involved in transcriptional regulation. ESC-specific regions are also closely located to genes involved in stem cell maintenance and development, while EpiLC-specific regions are associated with genes involved in differentiation (axon guidance and neural system development) (Figure S1C). Indeed, the largest change in probability is the increase in association of the term ‘multicellular organism development’ in EpiLC-specific regions. Taken together, these results suggests that OTX2 has different roles in naïve and formative pluripotency, acting near SOX motifs in ESCs at genes regulating stem cell maintenance and with ZIC proteins at genes that prepare for differentiation in EpiLCs.

### OTX2 modulates chromatin accessibility

Chromatin is remodelled during differentiation [23,24]. This enables cells to acquire different cell identities by opening new regulatory regions and by closing regions that regulate states that cells have transitioned beyond. As OTX2 has an essential role in orchestrating the choice between somatic and germline fates, we asked whether OTX2 functions to modulate chromatin accessibility. The assay for transposase-accessible chromatin with sequencing (ATAC-seq [25]) was used in wild-type and *Otx2^-/-^* ESCs, EpiLCs, and early during differentiation, as summarized in Figure 2A. To clearly distinguish between chromatin changes associated with early PGCLC differentiation and those associated with somatic cell differentiation, we compared *Otx2^-/-^* cells cultured in the presence of PGC-promoting cytokines with wild-type cells cultured in the absence of PGC-promoting cytokines. Under these conditions *Otx2^-/-^* cells produce an essentially pure (>90%) CD61^+^/SSEA1^+^ population [8,15,16] that we refer to as PGCLCs, while wild-type cells yield a cell population from which PGCLCs are absent (somatic cells) [8] (Figure S2). ATAC-seq data from the above differentiated samples was compared to ESCs and EpiLCs (Figure 2B). Principal component analysis (PCA) showed that loss of OTX2 does not dramatically alter the global chromatin accessibility in ESCs or EpiLCs, as wild-type and *Otx2^-/-^* cells cluster together at both stages. In contrast, after 2 days of differentiation, PGCLCs and somatic cells have drastically distinct chromatin accessibility landscapes (Figure 2B). Interestingly, d2 PGCLCs show an intermediate position between ESCs and EpiLCs suggesting that during differentiation into the germline, cells may restore some chromatin characteristics that previously defined ESCs. In contrast, wild-type differentiated cells (somatic cells) cluster far from other samples, suggesting that cells that adopt a somatic fate have drastic changes in chromatin accessibility compared to pluripotent cells and to PGCLCs (Figure 2B).

**Figure 2.**
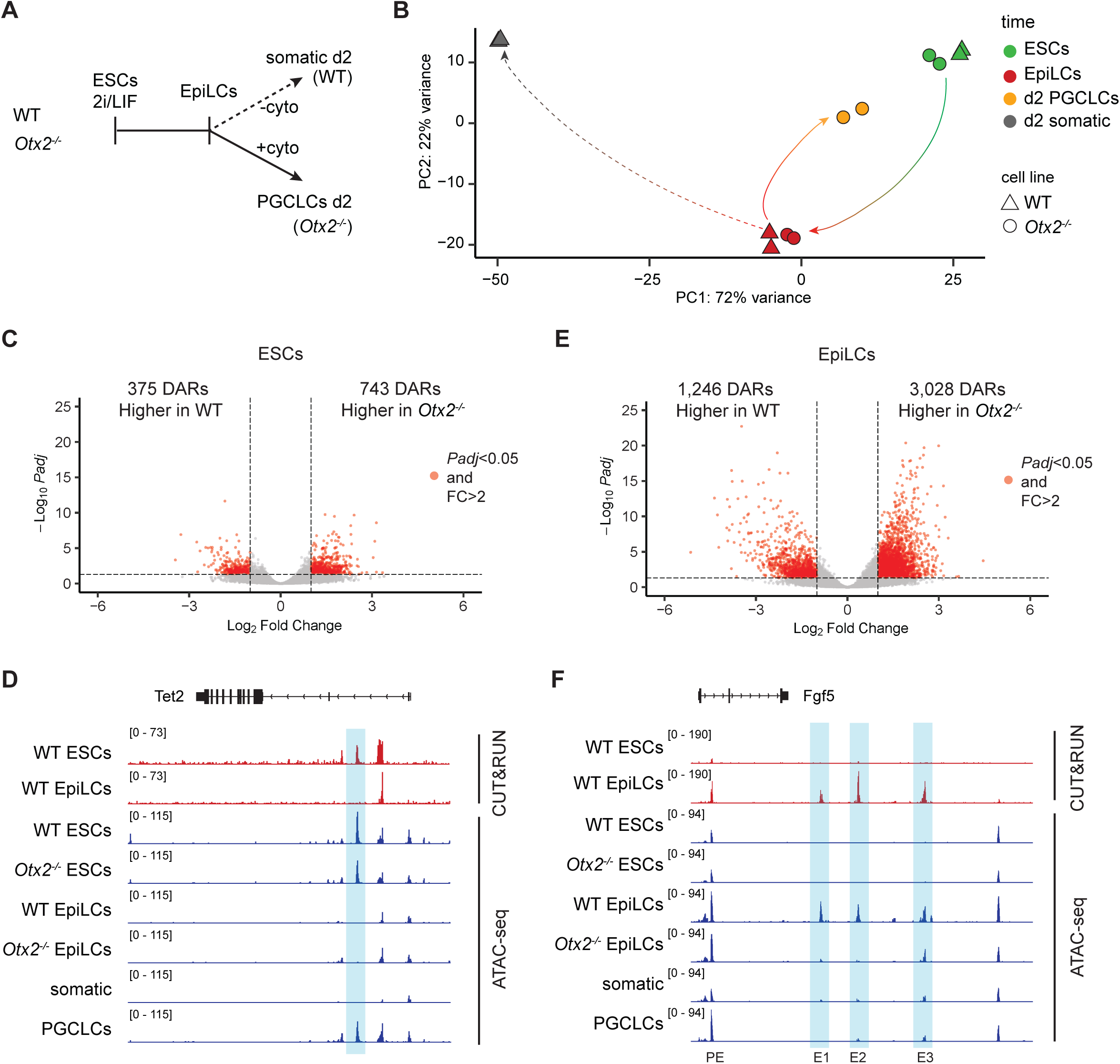
Chromatin accessibility during ESC - EpiLC - PGCLC/somatic transitions. A) Summary of ATAC-seq samples analyzed in this study. B) Principal component analysis of ATAC-seq samples. Arrows show ESC ◊ EpiLC, EpiLC ◊ PGCLC and EpiLC ◊ somatic cell transitions. C) Volcano plot comparing accessible regions in wild-type and *Otx2^-/-^* ESCs. D) OTX2 CUT&RUN and ATAC-seq tracks showing OTX2 binding (red) and accessibility (blue) at *Tet2*. DAR is highlighted in light blue. E) Volcano plot comparing differentially accessible regions in wild-type and *Otx2^-/-^*EpiLCs. F) OTX2 CUT&RUN and ATAC-seq tracks showing OTX2 binding (red) and accessibility (blue) at *Fgf5* locus. *Fgf5* enhancer nomenclature from [20]. DARs are highlighted in light blue.

Although the PCA suggests a high similarity between wild-type and *Otx2^-/-^* cells at both ESC and EpiLC stages, direct comparison shows that subsets of regions are differentially accessible. Comparing wild-type and *Otx2^-/-^* ESCs identified 375 differentially accessible regions (DARs) with increased accessibility in wild-type cells, and 743 regions with higher accessibility in *Otx2^-/-^* ESCs (Figures 2C). An example of ESC DARs where accessibility is increased in cells expressing OTX2 is the intragenic enhancer of *Tet2*. *Tet2* is expressed at high levels in ESCs but at low levels in EpiLCs [26]. Consistent with this changed expression, *Tet2* DAR has low accessibility and no OTX2 binding in wild-type EpiLCs (Figure 2D), suggesting that the T*et2*-associated enhancer becomes de-commissioned once cells leave the naïve pluripotent state. Interestingly, the *Tet2* DAR becomes accessible again in PGCLCs (Figure 2D).

Compared to ESCs, a higher number of DARs were detected from the analysis of EpiLCs. Specifically, 1,246 regions were more accessible in wild-type EpiLCs and 3,028 regions were more accessible in *Otx2^-/-^*EpiLCs (Figure 2E). An example of a gene with DARs that become accessible in EpiLCs is *Fgf5*. *Fgf5* becomes expressed after the exit from the naïve pluripotent state [28,29] Interestingly, *Fgf5* DARs correspond to characterized EpiLC-specific enhancers (PE, E1, E2 and E3)[20]. These enhancers either become accessible specifically in EpiLCs (E1, E2, E3) or show substantially more accessibility as ESCs transition to EpiLCs (Figure 2F). Moreover, the EpiLC specific DARs E1, E2 and E3 remain comparatively inaccessible in *Otx2^-/-^* EpiLCs (Figure 2F). This suggests that OTX2 acts as a pioneer TF [30] to regulate the accessibility of enhancers E1, E2 and E3.

### OTX2 induces accessibility in a subset of EpiLC regions

To investigate the hypothesis that changes in accessibility between wild-type and *Otx2^-/-^* cells are induced by OTX2, we analysed the overlap between regions that are more accessible in wild-type cells and OTX2 binding sites. In ESCs, OTX2 binds <10% (30 out of 375) of DARs that are more accessible in wild-type cells than in *Otx2^-/-^* cells (Figure 3A), suggesting that accessibility of ESC DARs is directly due to OTX2 in a small subset of DARs. In contrast, in EpiLCs, OTX2 binds almost 40% (446 out of 1,246) of the DARs that are more accessible in wild-type than in *Otx2^-/-^* cells (Figure 3B-C). Notably, these regions are mainly located distal to genes (91%, Figure 3D), despite the increased fraction of promoter regions bound by OTX2 in EpiLCs (Figure S1A). These 446 regions are more accessible in EpiLCs than ESCs, underscoring their EpiLC-specificity (Figure 3E). Without OTX2 these regions show the same low level of accessibility in both ESCs and EpiLCs in the absence of OTX2 suggesting that OTX2 may directly control accessibility at these sites (Figure 3E). This is borne out by motif analysis of these 446 regions, which shows a high enrichment for OTX2-like motifs (Figure 3F). Together, these results suggests that OTX2 is required to open these chromatin regions.

**Figure 3.**
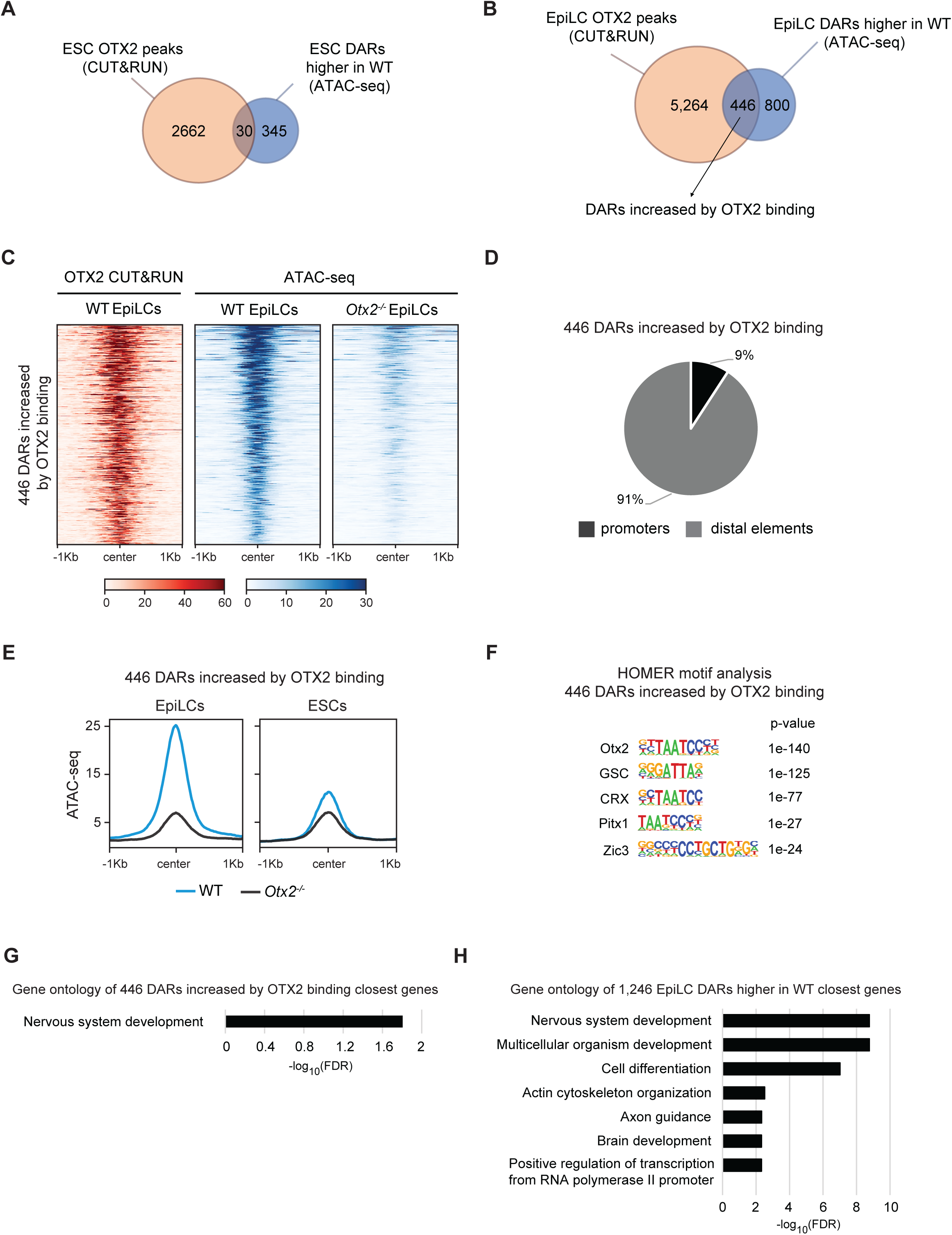
OTX2 facilitates chromatin accessibility in EpiLCs. A) Venn diagram of the overlap of OTX2-bound regions in ESCs (orange) and ESC DARs that are more accessible in wild-type ESCs (blue). B) Venn diagram of the overlap of OTX2 bound regions specific to EpiLCs (orange) and EpiLC-specific regions where accessibility is increased by OTX2 (blue) identifying 446 regions where OTX2 binding induces accessibility. C) Heatmap of OTX2 binding (CUT&RUN - red) and accessibility (ATAC-seq - blue) at 446 OTX2-bound regions that show increased accessibility in wild-type compared to *Otx2^-/-^* EpiLCs. D) Genomic distribution of the 446 OTX2-modulated regions. Promoter are +/- 1kb from any TSS. E) Read density profiles of ATAC-seq signal at the 446 OTX2-modulated regions in wild-type and *Otx2^-/-^*ESCs and EpiLCs. F) Motif analysis in 446 OTX2-modulated EpiLC regions showing high enrichment of OTX2-like motifs. G) Gene ontology analysis of the closest genes to the 446 DARs increased by OTX2 binding. H) Gene ontology analysis of the closest genes to the 1,246 EpiLC regions more accessible in the wild-type.

The remaining 800 DARs that are more accessible in wild-type EpiLCs than in *Otx2^-/-^* EpiLCs have an OTX2 CUT&RUN signal falling below the threshold applied in the bioinformatic analysis. However, CUT&RUN for OTX2 shows that these 800 DARs have low but detectable OTX2 binding (Figure S3A). Consistent with this, motif analysis of these 800 DARs identified OTX2 motifs, but with a reduced *p-value* compared to the 446 DARs mentioned above. Together with the increased *p-value* of ZIC motifs at these 800 DARs, this may indicate that these sites are bound by OTX2 at reduced affinity and that accessibility of these sites requires the combined action of OTX2 and ZIC TFs (Figure S3B). Gene ontology analysis of the genes closest to the 446 DARs increased by OTX2 binding reveals an association with nervous system development (Figure 3G). Expanding the analysis to the genes closest to the 1,246 DARs higher in the wild-type reveals that these regions are associated with terms of differentiation, specifically towards the neural fate (Figure 3H). These results suggests that OTX2 may instruct cells to differentiate towards the somatic fate by controlling accessibility at somatic regulatory regions.

### OTX2 expression in *Otx2*^-/-^ EpiLCs rescues chromatin accessibility

To determine whether OTX2 directly controls accessibility of the 1,246 regions that are more accessible in wild-type EpiLCs than in *Otx2*^-/-^ EpiLCs, we used an *Otx2^-/-^* cell lines carrying an OTX2-ER^T2^ fusion protein to rapidly induce OTX2. In these cells, OTX2-ER^T2^ is relocated from the cytoplasm to the nucleus within 20 minutes of tamoxifen addition [8]. To assess the effect of OTX2 on chromatin accessibility, *Otx2^-/-^*::Otx2-ER^T2^ ESCs were differentiated to EpiLCs and tamoxifen was added either one or six hours before the end of the differentiation (Figure 4A). ATAC-seq showed that a one-hour treatment with tamoxifen is sufficient to induce an increase in the average accessibility signal at these 1,246 regions (Figure 4B-C). The change in accessibility at one hour is almost exclusively due to the 446 regions that are bound by OTX2 strongly, as the remaining 800 regions showed little change in accessibility by one hour (Figure 4D). Although accessibility did increase further after six hours of tamoxifen treatment (Figure 4B-D), the fact that the major increase in accessibility occurred within one hour at the 446 regions strongly supports a direct role for OTX2 in opening these chromatin regions.

**Figure 4.**
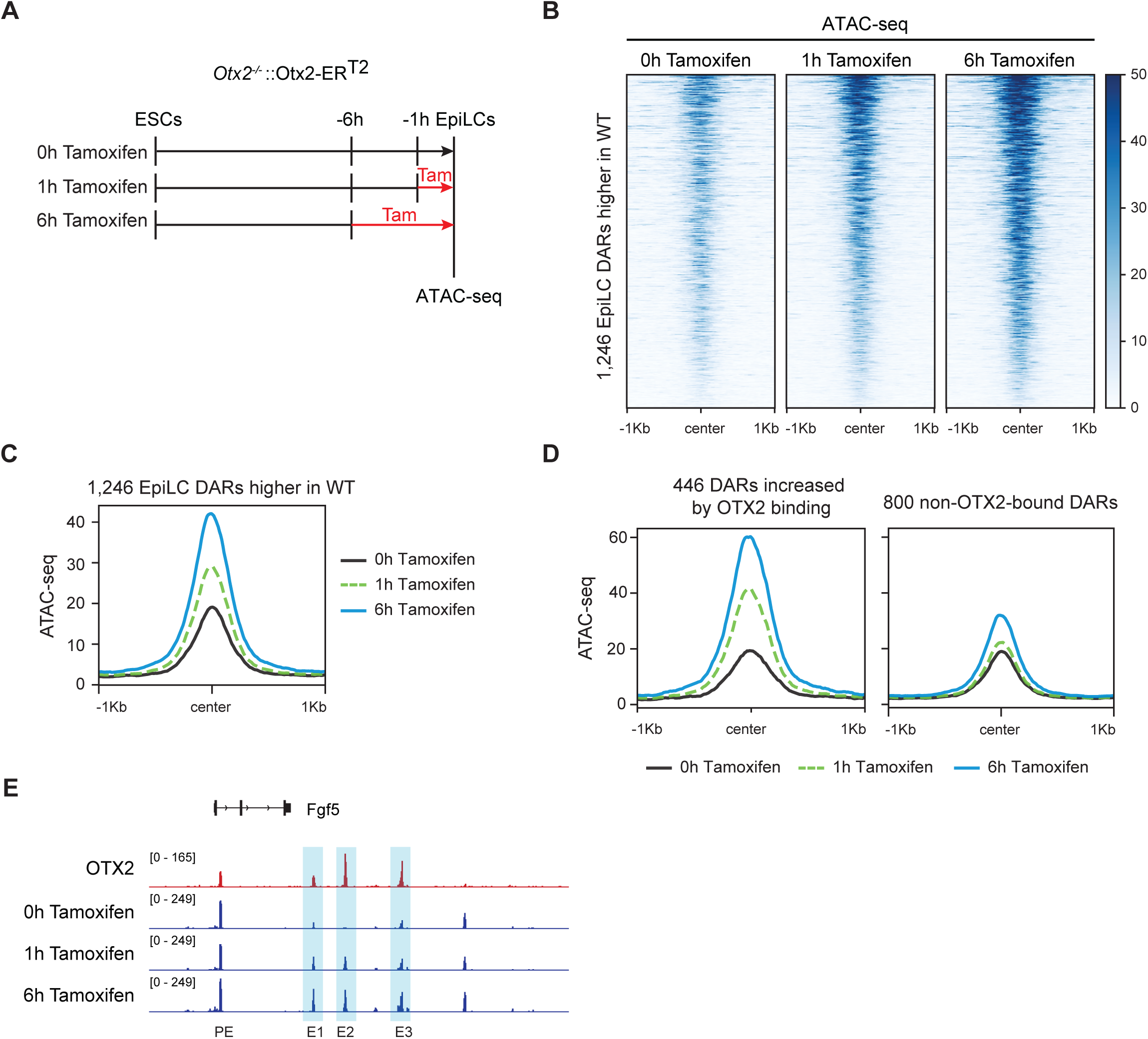
OTX2 expression rescues chromatin accessibility in EpiLCs. A). Differentiation scheme for *Otx2^-/-^*::Otx2-ER^T2^ ESCs into EpiLCs +/-Tamoxifen. B) Heatmap of ATAC-seq signal at the 1,246 EpiLC DARs higher in wild-type in *Otx2^-/-^*::Otx2-ER^T2^ EpiLCs treated for 1 or 6 hours with tamoxifen. C) Read density profile of ATAC-seq signal at the 1,246 EpiLC DARs higher in wild-type in *Otx2^-/-^*::Otx2-ER^T2^ EpiLCs treated for 1 hour (dashed green) or 6 hours (blue) with tamoxifen compared to untreated cells (black). D) Read density profile of ATAC-seq signal at the 446 DARs increased by OTX2 binding and the 800 non-OTX2-bound EpiLC DARs in *Otx2^-/-^*::Otx2-ER^T2^ EpiLCs treated for 1 hour (dashed green) or 6 hours (blue) with tamoxifen compared to untreated cells (black). E) OTX2 CUT&RUN and ATAC-seq tracks showing accessibility changes in response to OTX2 expression at the *Fgf5* locus. DARs are highlighted in light blue. *Fgf5* enhancer nomenclature from [20].

Among these OTX2-controlled regions are regulatory elements of the OTX2 target gene *Fgf5.* The accessibility of these regions increases after relocation of OTX2-ER^T2^; in the case of peak E2 from a baseline level (Figure 4E). This suggests that OTX2 facilitates *Fgf5* transcription not only by binding to enhancers but also by controlling the accessibility of these enhancers to the transcriptional machinery (Figure 4E). Together, these results suggest that OTX2 controls accessibility of a subset of putative enhancers in cells competent for both somatic and germline differentiation.

### OTX2 retains the ability to open chromatin during the early stages of germline differentiation

In wild-type cells, OTX2 restricts the number of cells entering the germline [8]. OTX2 can completely block entry of all cells into the germline when expression is enforced during the first 2 days of PGCLC differentiation [8]. To determine whether OTX2 retains the capacity to induce chromatin accessibility during PGCLC differentiation, *Otx2^-/-^*::Otx2-ER^T2^ EpiLCs were differentiated in the presence of PGC-inducing cytokines, either with or without tamoxifen. ATAC-seq and OTX2 CUT&RUN were performed at day 2 of differentiation (Figure 5A). Enforced OTX2 expression resulted in an increased accessibility at the 1,246 regions that were previously shown to be more accessible in wild-type EpiLCs than in *Otx2*^-/-^ EpiLCs (Figure 5B). These regions show low accessibility in the absence of tamoxifen but become open and bound by OTX2-ER^T2^ in tamoxifen treated cells (Figures 5B-C). This suggests that OTX2 can increase accessibility of chromatin regulatory regions not only in EpiLCs but also during the early stages of differentiation in the presence of PGC-inducing cytokines.

**Figure 5.**
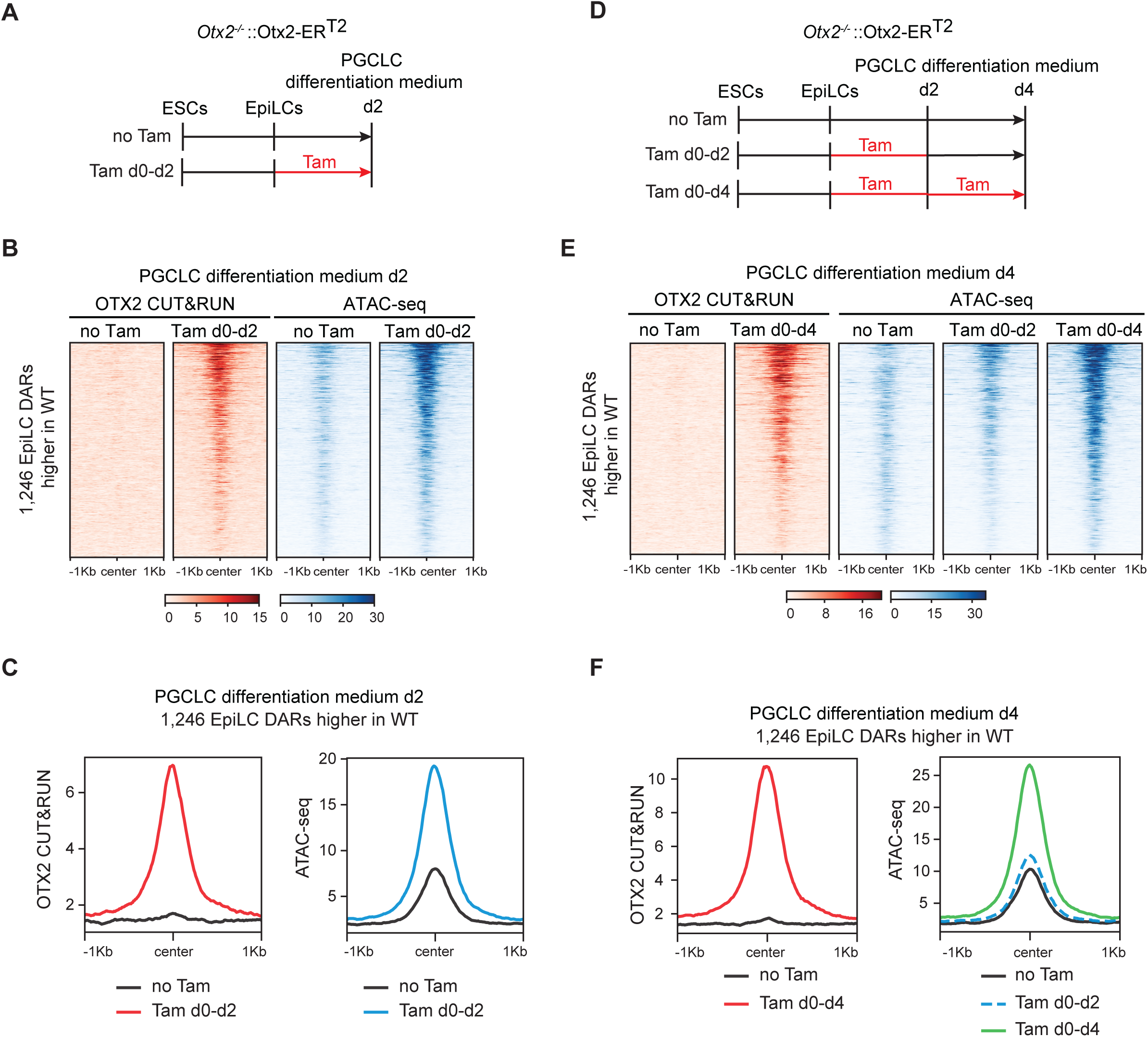
Sustained OTX2 expression in early PGCLCs induces chromatin accessibility. A) Differentiation scheme of *Otx2^-/-^*::Otx2-ER^T2^ ESCs to d2 PGCLCs +/-Tamoxifen. B) Heatmap of OTX2 CUT&RUN (red) and ATAC-seq (blue) signal in the 1,246 EpiLC DARs where accessibility is increased by OTX2 in Tamoxifen treated *Otx2^-/-^*::Otx2-ER^T2^ d2 PGCLCs. C) Read density profiles of OTX2 CUT&RUN (red) and ATAC-seq (blue) in day 2 PGCLCs +/-Tamoxifen at 1,246 DARs where accessibility is increased by OTX2. D) Differentiation scheme of *Otx2^-/-^*::Otx2-ER^T2^ ESCs to d4 PGCLCs +/-Tamoxifen. E) Heatmap of OTX2 CUT&RUN (red) and ATAC-seq (blue) in the 1,246 DARs where accessibility is increased by OTX2 in Tamoxifen treated *Otx2^-/-^*::Otx2-ER^T2^ d4 PGCLCs. F) Average profiles of OTX2 CUT&RUN (red) and ATAC-seq (blue) in day 4 PGCLCs +/-Tamoxifen at the 1,246 DARs where accessibility is increased by OTX2.

To determine whether OTX2 expression is essential to maintain chromatin accessibility in somatic cells, ATAC-seq was performed at day 4 of differentiation in PGCLC medium, in cells treated with tamoxifen for either the whole 4 days (Tam d0-d4) or just for the first 2 days (Tam d0-d2) (Figures 5D). The 1,246 DARs increased by OTX2 show higher accessibility at day 4 in cells treated throughout with tamoxifen (Figures 5E-F). In contrast, when tamoxifen is withdrawn after 2 days, these regions close by day 4 (Figures 5E-F). Together, these results show that OTX2 can induce accessibility in 1,246 chromatin regions in both EpiLCs and during differentiation in the presence of PGC-inducing cytokines. However, the continued presence of OTX2 is essential to maintain accessibility of these regions.

### Enforced OTX2 expression opens additional somatic regulatory regions

Since OTX2 can induce chromatin accessibility in EpiLCs, we speculated that OTX2 expression in somatic cells may also induce newly accessible regions. We compared tamoxifen treated *Otx2^-/-^*::Otx2-ER^T2^ d2 aggregates, which are blocked for PGCLC differentiation due to OTX2 activity, with untreated *Otx2^-/-^*::Otx2-ER^T2^ d2 aggregates (PGCLCs) and with wild-type ESCs and EpiLCs (Figure 2A). This identified 4,221 regions with high accessibility only in tamoxifen-induced *Otx2^-/-^*::Otx2-ER^T2^ d2 aggregates (Figure 6A). These regions are enriched for OTX2 motifs and to a lesser extent ZIC3 motifs (Figure 6B). CUT&RUN indicates that only a subset of the additional 4,221 accessible regions are bound by OTX2 (Figure 6C). Gene ontology analysis of the genes closest to these additional accessible regions reveals an association with cell differentiation and multicellular organism development, in particular neural system development (Figure 6D). Our results suggest that the enforced expression of OTX2 during the transition from EpiLCs to PGCLCs induces opening of additional somatic regions that may contribute to prevention of entry of cells into the germline.

**Figure 6.**
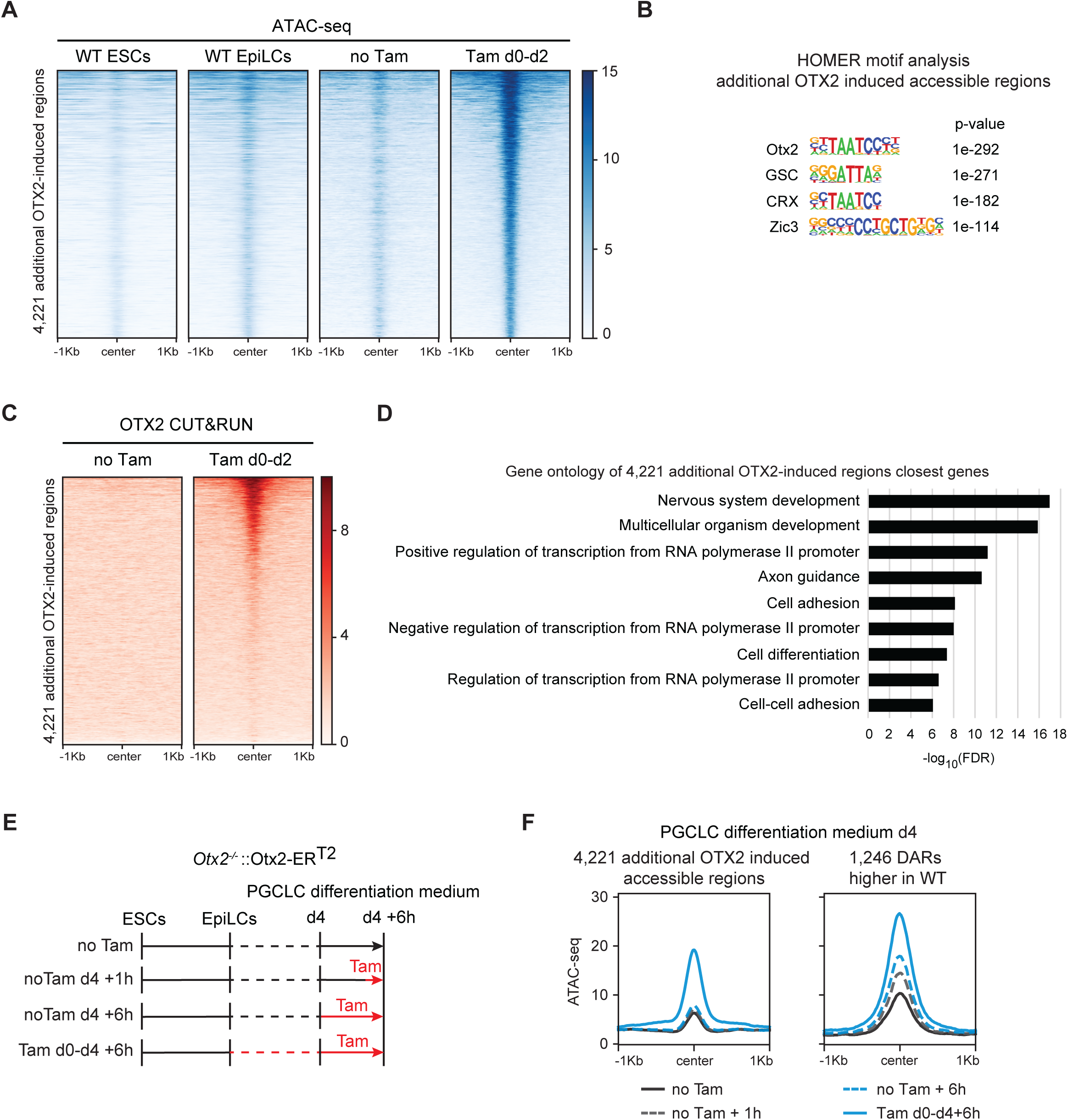
OTX2 induces accessibility at additional somatic regions. A) Heatmap of ATAC-seq signal in wild-type ESCs, wild-type EpiLCs and *Otx2^-/-^*::Otx2-ER^T2^ d2 PGCLCs treated +/-tamoxifen for 2 days at 4,221 additional OTX2-induced PGCLC regions. B) Motif analysis in the additional OTX2-induced PGCLC regions. C) Heatmap of OTX2 CUT&RUN signal in tamoxifen-treated *Otx2^-/-^*::Otx2-ER^T2^ d2 PGCLCs at 4,221 OTX2-induced PGCLC regions. D) Gene ontology analysis of the closest genes to the newly accessible PGCLC regions. E) Differentiation scheme of *Otx2^-/-^*::Otx2-ER^T2^ ESCs to day 4 PGCLCs with Tamoxifen treatments for 4 days + 6 hours, 6 hours and 1 hour only. F) Average profiles of ATAC-seq signal in *Otx2^-/-^*::Otx2-ER^T2^ PGCLCs treated with Tamoxifen as shown in Figure 6E at the 1,246 EpiLC DARs higher in WT and at 4,221 additional OTX2-induced PGCLC regions.

Since OTX2 is not able to block germline differentiation when relocated to the nucleus after day 2 of PGCLC differentiation [8], we tested the ability of OTX2 to open chromatin at both the 1,246 EpiLC regions and the 4,421 additional accessible regions. *Otx2^-/-^*::Otx2-ER^T2^ PGCLCs have been differentiated for 4 days in the absence of Tamoxifen and treated for one hour or six hours at day 4 (Figure 6E). While the 1,246 EpiLC regions show an increase in accessibility in response to Tamoxifen at day 4 in PGCLC differentiation medium, the 4,421 additional accessible regions are unable to respond to OTX2 and remain closed (Figure 6F). These results suggest that once cells have entered the germline, OTX2 can no longer modulate chromatin specifically at a subset of somatic associated regions.

## Discussion

At the implantation stage, cells within the mouse epiblast are required to undergo the important choice between somatic and germline differentiation. The transcription factor OTX2 plays a pivotal role in this choice, limiting the number of cells entering the germline, both *in vivo* and *in vitro*. In this work, we show how OTX2 works in cells competent for both somatic and germline differentiation to promote a somatic chromatin environment that primes the cells towards the somatic fate at the expense of the germline.

Using CUT&RUN we confirmed that OTX2 binds chromatin in both 2i/LIF ESCs and EpiLCs [20]. Motif analysis showed that these sites are, as expected, enriched for OTX2 binding motifs but also highlighted differences between the two cell types. In ESCs, sites bound by OTX2 are located distal to transcription units, are enriched for motifs of SOX and KLF TFs and are associated with genes involved in maintenance of the stem cell population. In EpiLCs, the majority of sites bound by OTX2 are also located distally, but a higher proportion are closer to promoter regions than in ESCs. This may suggest that OTX2 is more frequently involved in gene activation by binding to promoter regions as the cells exit naïve pluripotency. Moreover, in EpiLC, OTX2-binding sites are enriched for ZIC motifs, rather than the KLF and SOX motifs found in ESC sites. The absence of the KLF and SOX2 motifs in OTX2 EpiLC-specific binding site can be attributed to the decreased expression of both *Klf4* and *Sox2* mRNA after the exit from the naïve pluripotency [15,28,31,32]. These results indicate that while OTX2 is expressed in both ESCs and EpiLCs, the partner proteins that OTX2 acts alongside differ in naïve and formative pluripotent states.

The importance of protein partners in OTX2 function is also highlighted by the difference in the motif enrichment for the ZIC family of TF in the subsets of differentially accessible regions observed in EpiLCs. In the 446 regions where OTX2 binds strongly the only highly enriched motifs are OTX2-like. In contrast, in the remaining differentially accessible regions, OTX2 and ZIC motifs are equally enriched. *Zic2*, *Zic3* and *Zic5* are expressed in EpiLCs [31,33]. These two subsets of regions show different OTX2 occupancy, implying differences in OTX2 affinity between the 446 and 800 DARs. This suggests that OTX2 requires a co-activator to open the 800 regions. In epiblast stem cells (EpiSCs) that have lost competence for the germline, OTX2 works with ZIC2 to establish a new regulatory network that controls primed pluripotency [34,35]. Therefore, OTX2 may partner with ZIC2 or ZIC3 to control accessibility of somatic-associated regions that OTX2 cannot open on its own.

The ability of OTX2 to induce chromatin accessibility at somatic associated regulatory regions in cells that are competent for both somatic and germline fates suggests that the main action of OTX2 in the epiblast is to prime the cells to respond to somatic-directing cues and differentiate towards the somatic fate. As the somatic fate should be suppressed in developing PGCLCs to establish the PGC fate, somatic-associated regulatory regions should close. In this work, we showed that enforced expression of OTX2 during the transition to PGCLCs both maintains and induces opening of somatic-associated regulatory regions. OTX2 maintains its ability to open chromatin in only a subset of its regulated regions, as only regions that are already open in EpiLCs gain accessibility when OTX2 activity is induced in late PGCLCs. At this time, cells are already committed to germline differentiation and enforced expression of OTX2 cannot alter this cell fate. The additional regions where enforced expression of OTX2 induces accessibility during the initial 2 days of differentiation in the presence of PGC-inducing cytokines remain closed when OTX2 activity is induced at day 4 of PGCLC differentiation. This suggests, that genes regulated by these regions may induce and establish somatic identity.

In conclusion, in this work we generated novel information regarding the molecular mechanism of action of OTX2 in controlling the choice between somatic and germline differentiation. This shows that OTX2 can control accessibility of somatic-associated regulatory regions, preparing the cells to initiate the somatic fate at the expense of the germline. Regions that OTX2 opens in EpiLCs are always permissive to its action, while additional somatic associated regulatory regions become refractory to the presence of OTX2 once the germline fate is already established. Whether regulatory regions controlling genes that can block germline differentiation are among the latter subset is an interesting question for future analysis.

## Materials and Methods

### Cell culture

E14Tg2a [36], Otx2^-/-^ [2] and *Otx2^-/-^*::Otx2-ER^T2^ [8] cell lines were maintained in serum/LIF medium (Glasgow minimum essential medium (GMEM) Sigma, cat. G5154), 10% fetal bovine serum (Thermo Fisher Scientific, cat. 10270106), 2 mM L-glutamine (Invitrogen, cat. 25030-024), 1 mM pyruvate solution (Invitrogen, cat. 11360-039), 1× MEM non-essential amino acids (Invitrogen, cat. 11140-035), 0.1mM 2-Mercaptoethanol (Gibco, cat. 31350010), 100 U/ml homemade LIF in gelatin-coated flasks in a 37°C, 7.5% CO_2_ humidified incubator.

EpiLC and PGCLC differentiation was performed as previously described [16] with a few modifications. First, ESC grown in serum/LIF were dissociated with Trypsin (Thermo Fisher Scieitific, cat. 15090046), washed with PBS to remove any residual medium and resuspended in N2B27 medium (DMEM/F-12 (Thermo Fisher Scientific/Gibco cat. 21041025) supplemented with homemade N2 (DMEM/F-12, 11.1111 mg/ml apo-transferrin (Sigma, cat. T1147-100MG), 0.55% (w/v) Bovine Albumin Fraction V (Thermo Fisher Scientific, cat. 15260037), 2.2 μg/ml progesterone (Sigma, cat. P8783), 1.778 mg/ml putrescine (Sigma, cat. P5780), 3 mM sodium selenite (Sigma, cat. S5261) and 12.5 μg/ml insulin (Sigma, cat. I1882- 100MG) mixed 1:1 with Neurobasal (Thermo Fisher Scientific/Gibco cat. 12348017) supplemented with B27 without vitamin A (Thermo Fisher Scientific cat. 12587010), L-glutamine and penicillin-streptomycin (Thermo Fisher Scientific, cat. 15140122), followed by addition of 1.8 ml of 50 mM 2-Mercaptoethanol) supplemented with 0.4 μM PD0325901 (Stem Cell Technologies, cat. 72182), 3 μM CHIR99021 (APExBIO, cat. B5779) and 100 U/ml ESGRO LIF (Sigma/Millipore, cat. ESG1106) and adapted to 2i/LIF medium for at least 3 passages on poly-L-ornithine (Sigma, cat. P3655) and laminin (Corning, cat. 354232) coated 6-well plates. 1.0×10^5^ ESCs were washed with 1X PBS and plated on a well of 12-well plate pre-coated with 16.6 µl/ml fibronectin (Sigma, cat. FC010 and cat. F1141) in EpiLC medium: N2B27 medium supplemented with 20ng/ml Human Activin A (PeproTech, cat. 120-14), 12 ng/ml Human FGF-basic (Thermo Fisher Scientific, cat. 13256029) and 1% knock-out serum replacement (KOSR; Thermo Fisher Scientific/Gibco, cat. 10828028). Medium was freshly replaced after 24 hours. After 44 hours in EpiLC medium, cells were harvested with TrypLE Express Enzyme (Thermo Fisher Scientific, cat. 12604021), washed once with 1X PBS containing 1% Bovine Albumin followed by a second wash in 1X PBS. EpiLCs were then collected for analysis or for PGCLC differentiation.

1.5×10^5^ EpiLCs were resuspended in 5ml (3×10^4^ cells/ml) of GK15 medium (GMEM supplemented with 15% KOSR, 2mM L-glutamine, 1 mM pyruvate solution, 1× MEM non-essential amino acids, 100 U/ml penicillin/streptomycin, 0.1 mM 2-Mercaptoethanol) supplemented with 50 ng/mL bone morphogenetic protein (BMP) 4 (Qkine, cat. Qk038_BMP4_25 µg), 50 ng/mL BMP8a (R&D systems, cat. 1073-BP-010), 10 ng/mL stem cell factor (SCF) (R&D systems, cat. 455-MC-010), 10 ng/mL epidermal growth factor (EGF, R&D system, cat. 2028-EG-200), and 1,000 U/mL ESGRO and replated 100µl per well of a cell repellent U-bottom 96-well plate (Greiner Bio-one, cat. 650970). For cytokine-free differentiation, 1.5×10^5^ EpiLCs were resuspended in 5 ml of GK15 medium without cytokines. Cells were collected at day 2 and day 4 for analysis and day 6 for flow cytometry analysis.

*Otx2^-/-^*::Otx2-ER^T2^ cells were treated with 1 µM of 4-hydroxytamoxifen for the indicated times. To change tamoxifen containing media, aggregates were collected at day 2, washed, resuspended in GK15 and incubated in 5 mm dishes in rotation until day 4 and day 6 before collection and analysis.

### Flow cytometry

Flow cytometry analysis was performed as previously described [8,37] with minor changes. Cells were collected, dissociated using EDTA/Trypsin (Thermo Fisher Scientific, cat. 25200072) and neutralized in GK15 medium. Cells were centrifuged, washed with 1X PBS and resuspended in 100 µl 1X PBS/1% KOSR supplemented with Alexa Fluor 647 anti-CD15 (SSEA-1) (Biolegend, cat. 125607) and Phycoerythrin (PE) anti-CD61 (Biolegend, cat. 104307) antibodies diluted 1/200 and 1/600, respectively. Following a 10 min incubation at RT in the dark, cells were washed twice in 1X PBS and resuspended in 250 µl 1X PBS/1% KOSR supplemented with DAPI for live-cell selection. Acquisition was performed on a BD LSR Fortessa instrument and data was analysed using FlowJo v10.

### CUT&RUN

Cleavage Under Targets & Release Using Nuclease (CUT&RUN) was performed as previously reported [22] with no modifications using 5×10^5^ cells per replicate, 2 biological replicates per sample, 1:100 dilution of anti-OTX2 (R&D systems, cat. AF1979) and homemade pAG-MNase. Barcoded libraries were generated with the NEBNext Ultra II DNA Library prep kit (E7645S) and NEBNext Multiplex Oligos for Illumina (E7335S, E7500S, E7710S, E7730S) with modifications. 5 ng of DNA was used per library. End-prep and ligation were performed according to the manufacturer’s instructions. 1 µl of USER enzyme was used per sample. For the final PCR step, 1 µl of both universal and indexed primers were used, and the annealing/extension step was decreased to 30 seconds to reduced amplification of large fragments. Clean up steps were performed with home-made PCR purification beads. Library quality was assessed with HSD1000 ScreenTapes (Agilent, cat. 5067-5584 and 5067-5585) in a Tapestation 2200 (Agilent). Libraries were sequenced paired-end on Illumina NextSeq 500 and NextSeq 2000.

### CUT&RUN analysis

Quality of reads was assessed by FastQC (v. 0.11.9 [38]), index and adaptor sequences were trimmed using TrimGalore (v. 0.6.6 [39]) and Cutadapt (v. 1.9.1 [40]) followed by a second quality check with FastQC. Sequences were aligned to the mm10 mouse reference genome using Burrows-Wheeler Alignment (BWA, v. 0.7.16 [41]) tool with the maximal exact matches (MEM) option. Fragments were quality filtered (>10) with Samtools (v. 1.6[42]) and PCR duplicates were marked with Picard (v. 2.23.3 [43]) and removed with Samtools. Reads of OTX2 CUT&RUN in ESCs and EpiLCs were normalized to the number of e.coli reads before further analysis.

Peaks were called with Model-Based Analysis of ChIP-Seq (MACS2, v. 2.1.1 [44]) at 5% FDR using parameter-f BAMPE for paired-end input. Blacklisted regions were subtracted from the peak list using BEDTools (v. 2.27.1 [45]) and reproducible peaks between the 2 biological replicates were identified with IDR (v. 2.0.4.2 [46]). Bedtools was used to identify ESC-specific, EpiLC-specific and common OTX2 bound regions and following intersections with ATAC-seq data. Bigwig tracks were generated with the bamCoverage option in the python-based DeepTools (v. 3.5.1 [47]).

### ATAC-seq

ATAC-seq was performed with modification from the original Buenrostro protocol [25,48] using 5×10^4^ cells per replicate, 2 biological replicates per sample. Cells were washed in PBS, incubated for 20 minutes in lysis buffer (10 mM Tris-HCl pH 7.4, 10 mM NaCl, 3 mM MgCl_2_, 0.1% (v/v) Igepal) on ice. After centrifugation, nuclei were resuspended in 25 µl of freshly prepared Tagmentation Master Mix containing Assembled Trasposomes (Active Motif, cat.

53150). Tagmentation was performed in a thermocycler at 37°C for 20 minutes. Tagmented DNA was recovered using the DNA Clean and Concentration-5 kit (Zymo Research, cat. D4013). Library was prepared using NEBNext High Fidelity 2x PCR Master Mix and indexed primers reported in [25] using the following protocol: 72°C for 5 minutes, and 98°C for 30 seconds followed by 9 cycles of: 98°C for 10 seconds, 63°C for 30 seconds, 72°C for 1 minute. Library quality was assessed with HSD1000 tapes in a Tapestation Instrument. Libraries were sequenced paired-end on Illumina NovaSeq 6000 and NextSeq 2000.

### ATAC-seq analysis

ATAC-seq analysis was performed with a pipeline similar to CUT&RUN with the following modifications. After alignment with BWA, mitochondrial fragments were eliminated with Samtools. Mapped reads were shifted by +4 bp for the forward strand and-5 bp for the reverse strand. ESCs, EpiLCs, PGCLC d2 and somatic cells were normalized by read depth before peak calling. Shifted bam files were transformed into BED files for peak calling using MACS2.

A list of all accessible regions in the 6 samples analysed in this study was created and used as input for multiCov tool from the deepTools suite together with bam files to retrieve the coverage of each region in each sample. The resulting matrix was used as input for differential accessibility analysis pairwise comparisons with DESeq2 (v. 3.18 [49]). Volcano plots were generated by DESeq2.

### Peak annotation

Lists of identified regions have been analysed with R package ChIPseeker (v 3.8 [50,51]) to annotate their position corresponding to the closest gene. The results of ChIPseeker were used to generate pie charts. The list of closest genes was used as input for Gene Ontology analysis.

### Motif and Gene Ontology analyses

Motif analysis was performed with Hypergeometric Optimization of Motif EnRichment (HOMER v. 4.11 [52]) using findMotifsGenome.pl and options-size given-nomotif. Gene ontology was performed with Database for Annotation, Visualization and Integrated Discovery (DAVID) database [53,54] using the ENSEMBL gene annotation for the closest gene to each region annotated by ChIPSeeker.

## Data visualization

Average bigwigs between 2 biological replicates were generated with the deepTools suite (bigwigAverage). The average bigwig file was used to generate heatmaps and average profiles with computeMatrix (reference-point, options--referencePoint center--sortRegions descend--skipZeros--missingDataAsZero), plotHeatmap and plotProfiles of the deepTools suite.

## Data availability

Original ATAC-seq and CUT&RUN data are deposited at the NCBI Gene Expression Omnibus (GEO) with accession numbers GSE289298 (ATAC-seq) and GSE289297 (CUT&RUN).

## Acknowledgements

We thank Claire Cryer and Fiona Rossi for assistance with flow cytometry, Edward Foo for preliminary ATAC-seq analysis, the Genetics Core at the Western General for NGS sequencing and all members of the Chambers lab for feedback and suggestions. pAG-MNase was a gift from Abdenour Soufi and Burak Ozkan (University of Edinburgh). This work was funded by UK Medical Research Council Grant MR/T003162/1 and by a Marie Sklodowska-Curie fellowship (H2020-MSCAIF-2018/843879) to EB.

## Author contributions

EB and IC conceived the project. EB designed, performed and analysed experiments. EB and IC wrote the manuscript.

---

**Supplementary Figure S1.**
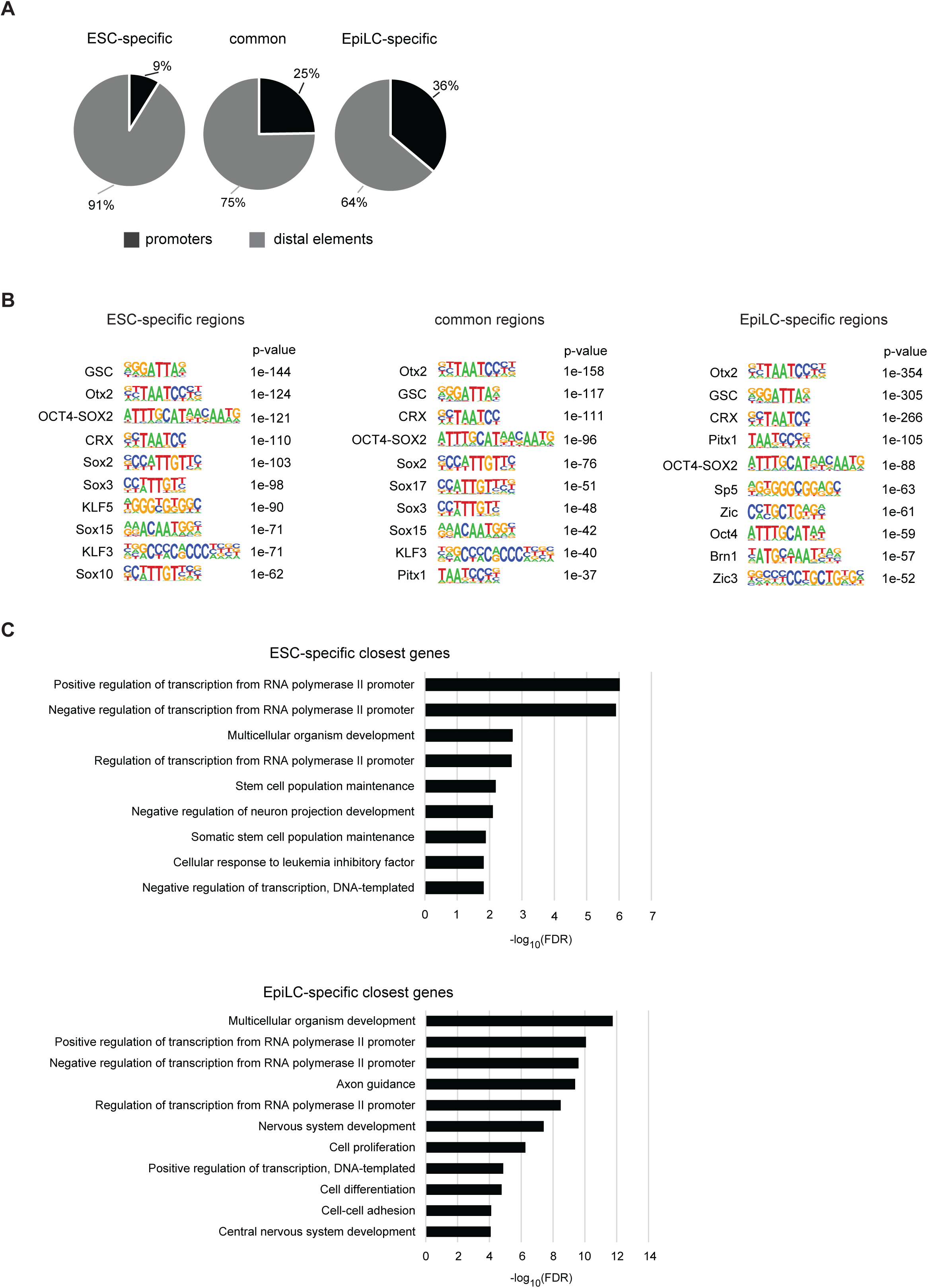
A) Genomic distribution of OTX2 bound regions; promoters are defined as +/- 1kb from a TSS. B) Motif analysis in ESC-specific, common and EpiLC-specific regions. C) Gene ontology of the closest genes to ESC-specific and EpiLC-specific OTX2 bound regions.

**Supplementary Figure S2.**
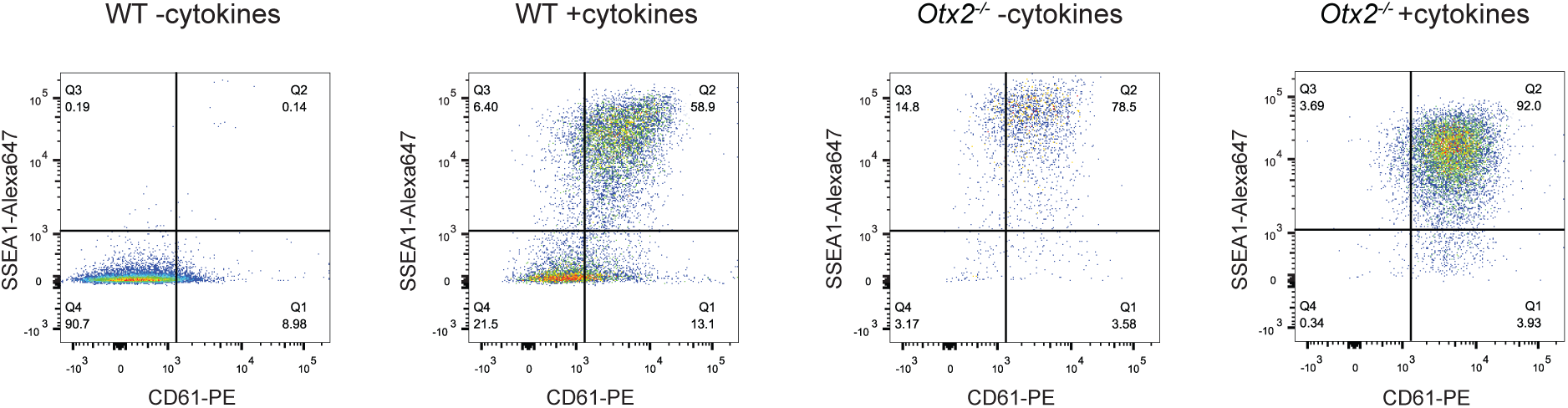
Flow cytometry plots showing surface expression of CD61 and SSEA1 in d6 aggregates from wild-type and *Otx2^-/-^* cells grown in the presence or absence of PGCLC inducing cytokines.

**Supplementary Figure S3.**
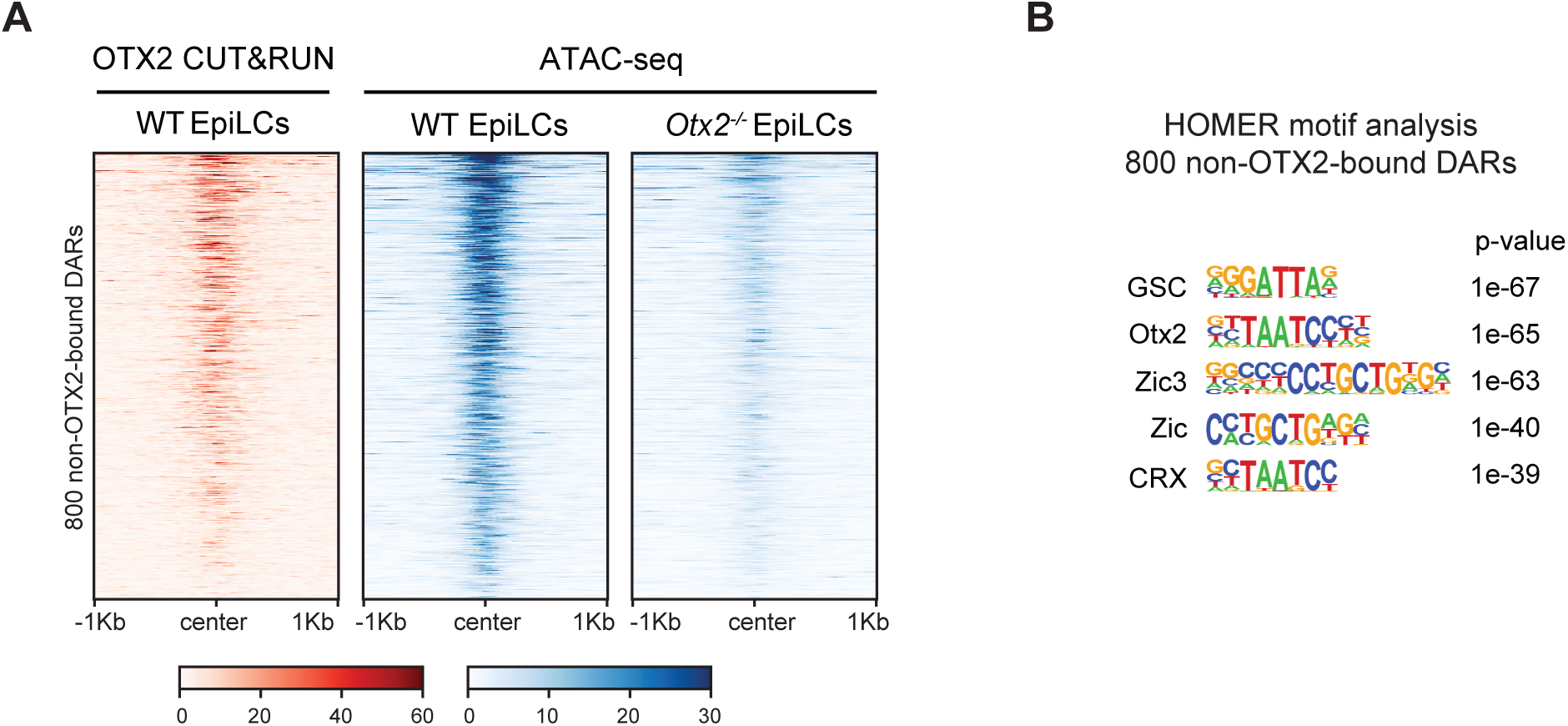
A) Heatmap of OTX2 CUT&RUN (red) and ATAC-seq (blue) signal in 800 non-OTX2-bound EpiLC DARs. B) Motif analysis in 800 non-OTX2-bound EpiLC DARs showing high enrichment of motifs recognised by both OTX2 and ZIC family members.

